# Recombination Disequilibrium in Ideal and Natural Populations

**DOI:** 10.1101/035121

**Authors:** Yuan-De Tan

**Author notes:** Corresponding author: Yuan-De Tan.

## Abstract

Following Hardy-Weinberg disequilibrium (HWD) occurring at a single locus and linkage disequilibrium (LD) between two loci in generations, we here proposed the third genetic disequilibrium in population: recombination disequilibrium (RD). RD is a measurement of crossover interference among multiple loci in a random mating population. In natural populations besides recombination interference, RD may also be due to selection, mutation, gene conversion, drift and/or migration. Therefore, similarly to LD, RD will also reflect the history of natural selection and mutation. In breeding populations, RD purely results from recombination interference and hence can be used to build or evaluate and correct a linkage map. Several practical examples from F_2_, testcross and human populations indeed demonstrate that RD is useful for measuring recombination interference between two short intervals and evaluating linkage maps. As with LD, RD will be important for studying genetic mapping, association of haplotypes with disease, plant breading and population history.

---

In the early days of the last century, after rediscovery of Mendel’s genetic laws, one began to look at the genetic behavior of a pair of alleles at a single locus in an ideal population. As a result, Hardy and Weinberg each independently discovered an equilibrium law of frequencies of both alleles and genotypes at a single locus in a very large population in 1908 (Hardy 1908; Weinberg 1908). This equilibrium law, called Hardy and Weinberg equilibrium (HWE), is a foundation of population and evolutionary genetics and an important landmark in population genetics. Now it is a common practice to check whether the observed genotypes conform to HWE expectations in disease gene mapping. These expectations appear to hold for most human populations but at some particular marker sites deviation of allele frequencies from HWE may occur, suggesting that the problems are associated with genotyping or population structure or, in samples of affected individuals, the markers are associated with disease susceptibility (Wigginton *et al*. 2005).

When, however, our interest was extended from one-locus system to two-locus system where each locus has a pair of alleles, a very important phenomenon, linkage disequilibrium (LD), was uncovered (Geiringer 1944; Hill and Robertson 1968; Lewontin and Kojiana 1960; Lewontin 1964; Robbins 1918). LD plays a fundamental role in gene fine mapping and especially in study of genome-wide association of genetic variants and diseases of interest. Patterns of LD have become a tool for fine mapping of genes for a complex disease of study (Hastings 1984; Pritchard and Przeworski 2001) and are a topic of a great current interest due to single nucleotide polymorphisms (SNPs). LD is also of interest for what it can reveal the evolution of population because the patterns of LD are determined, in part, by population history. LD throughout genome reflects the population history and the pattern of geographic subdivision, whereas in a genomic region LD reflects the history of natural selection, gene conversion, mutation (Slatkin 2008).

What happens when a three-locus system is considered? Several studies attempted to generalize LD from two-locus system to three-locus system. For example, Bennett(Bennett 1954) defined three-locus LD as

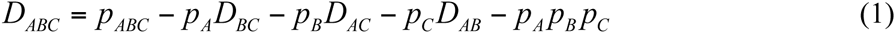

where *p_ABC_* is the frequency of gamete ABC, *p_A_*, *p_B_*, *p_C_* are frequencies of alleles A at locus *a*, B at locus *b*, and C at locus c, respectively, and *D_ij_* = *p_ij_* − *p_i_p_j_* is the standard two-locus LD definition where *i* = *A*, *B*, *j* =*B*, *C* and *i* ≠ *j*. But these attempts have not had substantial progress in the past several decades because LD at three-locus level or higher level becomes very complicated and so poor application has also made in practice due to recombination interference. Hastings (Hastings 1984) indicated that commonly used measures of linkage disequilibrium are not appropriate for a multilocus setting. Thomson and Baur (Thomson and Baur 1984) also showed by an example that combinations of allele frequencies and pairwise linkage disequilibrium terms, each of which is permissible at the two-locus level, may not be permissible at three-locus level. Actually, in two-locus system the significance of linkage disequilibrium in biology and evolution is quite obvious, while in three-locus or multilocus system, double or multiple crossovers and crossover interference become a major component of linkage. Recall crossover interference occurring in three-locus system. In classical genetics, crossover interference is measured by coefficient of coincidence defined as 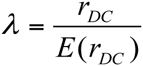 where *r_DC_* and *E*(*r_DC_*) are the observed and expected frequencies of double crossover types between two neighboring intervals *a-b* and *b-c*, respectively, where the locus order is assumed to be *a-b-c*. *E*(*r_DC_*) = *r_ab_r_bc_* under condition of no crossover interference where *r_ab_* and *r_bc_* are the observed recombination fractions between loci *a* and *b* and between loci *b* and *c*, respectively. Coefficient of coincidence as a measurement of crossover interference is due to the fact that only the positive crossover interference has been discovered in early genetic study. With the advance of technologies in molecular genetics, in particular, with the wide application of genotyping at molecular markers such as microsatellites and single nucleotide polymorphisms (SNPs), negative interference has widely been discovered. The coefficient of coincidence is not appropriate to describe negative interference because, for example, for positive interference, 0 ≤ *r_DC_* ≤ *r_ab_r_bc_* so that 0 ≤ *λ* ≤ 1, but for negative interference, *r_DC_* ≥ *r_ab_r_bc_* ≥ 0, then *λ* ≥ 1. Thus the coefficient of coincidence *λ* is asymmetric for interference in the positive and negative directions, which leads to difficulty of testing for the positive or negative interference in statistics. For example, Esch (Esch 2005) trialed to adopt null simulation way instead of direct way to test coefficient of coincidences. However, simulation has a big defect that the deviation of coefficient of coincidence from expected value *E*(*λ*) = 1 (no interference) would significantly increase with decreasing distance between loci (Esch 2005). So null simulation also is not a good way. However, both the positive and negative interferences in effect cause recombination disequilibrium (RD). RD is defined as difference (*D_r_*) between the observed and expected frequencies of double or multiple crossover types in a population. Obviously RD is symmetric in both negative and positive interference directions. It will be easy to establish statistics for RD test. In this paper, we propose a mathematical definition of RD and a statistical method for hypothesis test of RD in ideal and natural populations and then we give some practical applications.

To estimate and test for RD among loci, it is required to estimate the three-locus gamete frequencies. In a natural population, especially, in a human population, for codominant markers, many existing powerful methods such as PHASE (Stephens *et al*. 2001), fastPHASE (Scheet and Stephens 2006), BEAGLE (Browning and Browning 2007), IMPUTE2 (Howie *et al*. 2009) and MaCH (Li *et al*. 2010) can be used to estimate haplotype frequencies. In F_2_ population, also a lot of expectation maximization (EM) algorithms (Esch 2005; Li *et al*. 2007; Sergeev and Arapova 2002) can be used to estimate triplet gametic frequencies way can be utilized to distinguish dominant homozygous genotypes from heterozygous genotypes (Esch 2005; Liu 1998; Sergeev and Arapova 2002), the EM algorithms have low power to estimate gametic frequencies in F_2_ population. Tan and Fu (Tan and Fu 2007) to develop a new approach (TF) to promote estimation accuracy of three-locus dominant gamete frequencies in F_2_ population. Different from the current EM methods(Esch 2005; Li *et al*. 2007; Sergeev and Arapova 2002), the TF method is based on such an assumption that sister-gametes have the same frequencies in F2 population, which greatly reduce complexity of mathematical equations so that the power is significantly promoted. This assumption is correct because as indicated above, F_2_ population is ideal population without mutation, selection, genetic drift and immigration. In our current work, we use a modified TF method (see Supplementary Notes S1 and S3 for detail) to estimate dominant triplet gametic frequencies and a very fast and highly accurate method to estimate frequencies of triplet codominant gametes in an F_2_ population (see Supplementary Notes 2 and S3) because their algorithms are too trivial and too complex and have no open source software package to be used. In this paper we focus on RD instead of estimation of gametic frequencies.

## Definition And Tests

Since these four *q*’s are estimated separately, sum of them does not always satisfy a constraint of *q* = *q*_1_ + *q*_2_ + *q*_3_ + *q*_4_ = 0.5. For this reason, we constrain sum of estimates to 0.5:

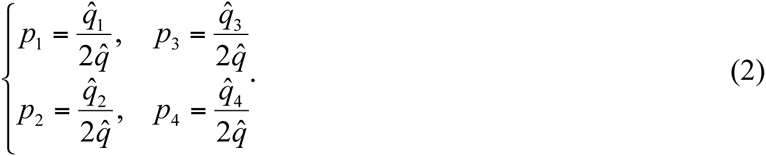

For linked loci, frequencies of the four pairs of nonsister gametes can be used to distinguish coupling phase from repulsion phase between loci, which allows us to find double crossover types. For example, suppose an order of three loci is a-b-c. Then if *p*_4_ is smallest and *p*_1_ is largest, each allele pair is in coupling phase and *p*_4_ is frequency of double crossover types; if *p*_4_ is largest and *p*_1_ is smallest, then allele A at locus *a* and allele C at locus *c* are in coupling phase but allele B at locus *b* is in repulsion phase, so *p*_1_ is frequency of double crossover types. On the other hand, if *p*_2_ is largest and *p*_3_ is smallest, then allele A at locus *a* and allele B at locus *b* are in coupling phase but allele C at locus *c* is in repulsion phase, then *p*_3_ is frequency of double crossover types. Similarly, if *p*_2_ is smallest and *p*_3_ is largest, then *p*_2_ is frequency of double crossover types. Thus the smallest *p* must be frequency of double crossover types. When, however, negative inference occurs, the case would be different from the above: the gamete with the smallest frequency is not necessarily double crossover types because negative interference increases frequency of double crossover types in a population. But the law that the gamete with the largest frequency in a population is parental type still holds because negative interference occurs only when one or both two neighboring intervals are very short.

In coupling phase *p*_4_ is frequency of double crossover types. Thus, recombination fractions between loci *a* and *b*, between loci *b* and *c*, and between loci *a* and *c* can be calculated by

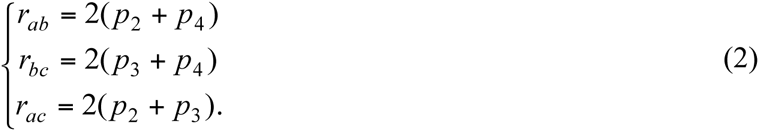

Recombination fractions between loci in the other orders in coupling phase are also given in a similar way.

In repulsion phase, the order (*a-b-c*) determines *p*_1_ to be frequency of double crossover types, thus the recombination fractions between loci *a* and *b*, between loci *b* and *c*, and between loci *a* and *c* are

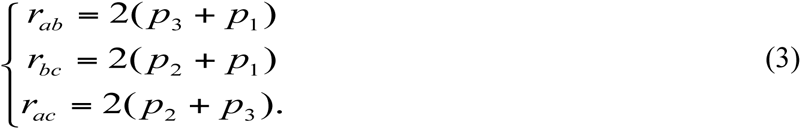

The recombination fractions between loci in the other orders in repulsion phase can be estimated in a similar fashion.

Thus, if, in coupling phase, *p*_4_ is frequency of double crossover types, RD is defined as difference between the observed and expected frequencies of double crossover types, that is,

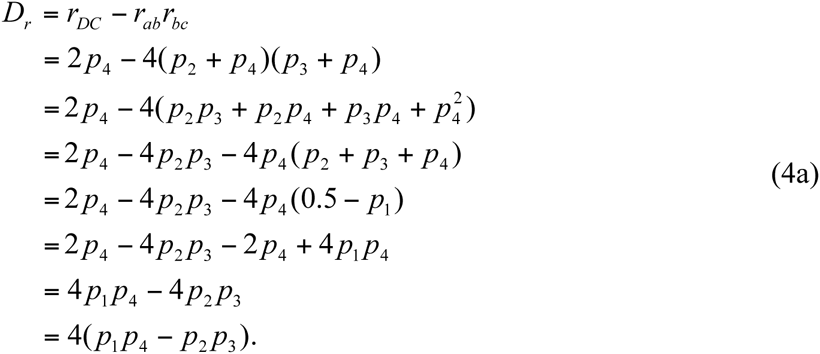

in repulsion phase, we also have

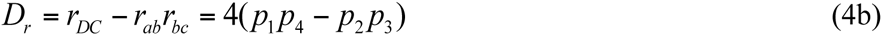

if frequency of double crossover types is *p*_1_, or

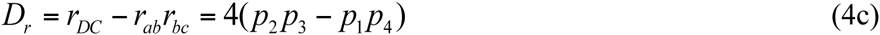

if *p*_2_ or *p*_3_ is frequency of double crossover types. Obviously, *D_r_* is a symmetric measurement of positive and negative interferences: *D_r_* = 0 means that recombination between two neighboring intervals is in equilibrium, say, recombination events occur independently between these two neighboring intervals. Negative recombination interference occurs if *D_r_* > 0 or positive recombination interference occurs if *D_r_* < 0. The maximum RD is *D_r_* = 4 *p*_1_*p*_4_ and the minimum RD is *D_r_* = −4*p*_2_*p*_3_ in systems AbC/aBc and ABC/abc or the maximum RD is *D_r_* = 4*p*_2_*p*_3_ and the minimum RD is *D_r_* = −4*p*_1_*p*_4_ in systems aBC/Abc and ABc/abC. Both positive and negative recombination interferences occur in two very short intervals of chromosomes. But positive interference may be physical interference due to rigidity of DNA chains while negative interference may be biological interference due to activation of enzymes. One can predict that activation of enzymes on promoting recombination of DNA would become weak or disappear as interval distances increase. Similarly to the LD test, a standard chi-square statistic can be used to test for RD among three loci. In the ABC/abc system, the chi-square statistic is

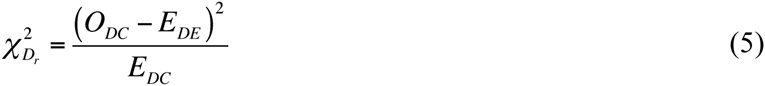

where *O_DC_* and *E_DC_* are numbers of observed and expected double crossover types and *N* is number of individuals sampled from F_2_ population: *O_DC_* can be given by using frequency of double crossover types and sample size N: *O_DC_* = *Nr_DC_* and *E_DC_* can be obtained from expected frequency of double crossover types and N: 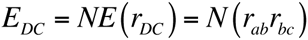. Inputting them into Equation (5), we have

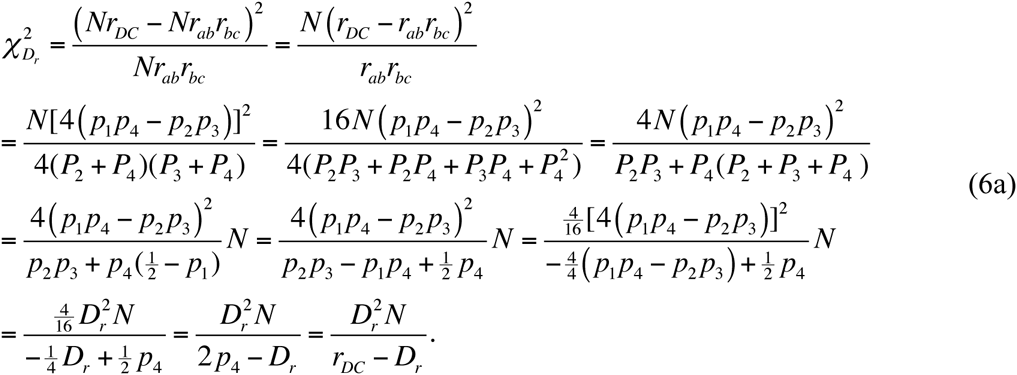

In the AbC/aBc system, the chi-square statistic for RD is

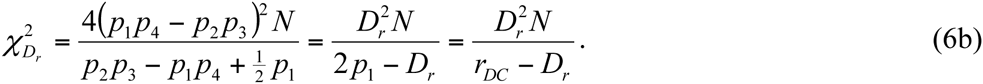

In the aBC/Abc system, the chi-square statistic for RD is

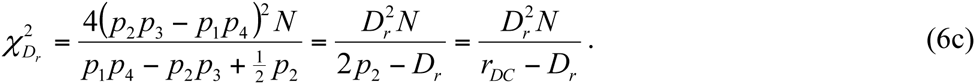

And in the ABc/abC system, the chi-square statistic for RD is

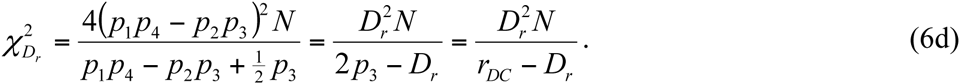

For codominant markers, we will convert AbC/aBc and ABC/abc to 101/010 and 111/000. Similar way can also be used to the other genotypes. Under the null hypothesis H_0_, these four chi-square statistics have an approximate *χ*^2^ distribution with 1 degree of freedom.

Theoretically, RD reflects recombination disequilibrium due to interference in an ideal population (such as F_2_ or backcross population). In a natural population, RD could result from evolutionary factors such as selection, migration, mutation, drift, and gene conversion because these factors change allele and gamete frequencies which may make frequencies of sister gametes unequal. For the case of *p*(*ABC*) ≠ *p*(*abc*), *p*(*Abc*) ≠ *p*(*aBC*), *p*(*ABc*) ≠ *p*(*abC*), or *p*(*AbC*) ≠ *p*(*aBc*), the RD can be given by

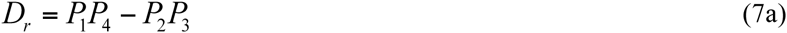

in coupling phase or

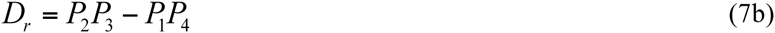

in repulse phase where

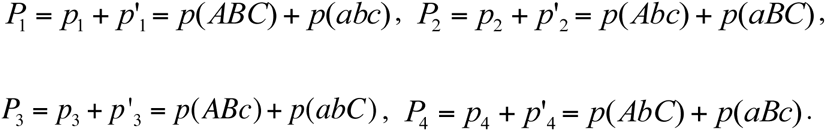

Derivation of Equation (7) is given in Supplementary Note S3. Chi-square test for RD among loci in natural population has the same expression with Equation (6). Supplementary Note S4 gives derivation of Equation (6a) in the case of which sister gametes have unequal frequencies. In an ideal population, Equation (B6) is reduced to Equation (4), underlying that if all three loci follow Hard-Weinberg equilibrium and sister gametes have equal frequencies, their RD derives only from recombination interference, otherwise, the RD results from selection, mutation, gene conversion, migration or drift and/or recombination interference.

## APPLICATIONS

### Test for recombination disequilibrium among dominant and codominant molecular markers in F_2_ population

As an example for illustrating construction of linkage map by MAPMAKER/EXP (version 3.0b), LANDER *et al*. (Lander *et al*. 1987) provided a RFLP dataset of 333 F_2_ mice. Since RFLP markers are codominant in the genotype dataset, uppercase letters A, H, and B are used to denote homozygote A (a pair of the same alleles coming from parent A), heterozygote H, and homozygote B (a pair of the same alleles coming from parent B) at a locus, respectively. We applied our moment method (Supplementary Note S2) to the original dataset of the first 6 codominant markers and the modified TF (Supplementary Note S1) to dominant marker data converted by changing B to H to estimate frequencies of four nonsister gametes. Using these gamete frequencies, we calculated and tested for these 20 triplet RDs. The results are summarized in Tables 1~3. Table 1 displays the frequencies of the four nonsister gametes estimated by TF in 333 F_2_ mouse individuals. One can find that RDs in 7 triplets (123), (125), (126), (135), (145), (235), and (245) are significant where 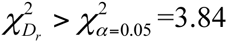 at *p*- *value* < 0.05. However, for the same genotype data, Table 2 obtained by performing the modified TF method shows that only triplets (123) and (145) have 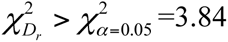. As codominant markers can display the full linkage information for estimation of gamete frequencies in F_2_ population and have no phase problem(Liu 1998), the moment estimates of gamete frequencies in the codominant three-locus systems are pretty accurate (Table 3), which make RD estimation more reliable. The result obtained from codominant data shows that only triplet (123) had a significant RD. These indicate that estimate of gametic frequencies in a dominant three-locus system in F_2_ population provided by the modified TF method are very close to the estimates of gametic frequencies in codominant three-locus system. The three methods all found that triplet (123) had a significant positive RD, therefore, we can make a conclusion that there was a significant negative crossover interference among loci 1(T175), 2 (C35), and 3(T24). This conclusion is well validated by mapping of these three markers into two short intervals (T175-C35 in 3.1 cM and C35-T93 in 13.5cM (Tan and Fu 2007)).

**Table 1.** The moment estimated frequencies of nonsister gametesin triplets of 6 dominant loci in 333 F2 mice^a^ and recombination disequilibrium tests

| Locus | | | $p_1 =$ | $p_2 =$ | $p_3 =$ | $p_4 =$ | RD | |
| --- | --- | --- | --- | --- | --- | --- | --- | --- |
| a | b | c | $p(abc)$ | $p(Abc)$ | $p(abC)$ | $p(aBc)$ | $D_r$ | $\chi^2$ |
| 1 | 2 | 3 | 0.200547 | 0.089443 | 0.097526 | 0.112484 | 0.055341 | 5.99423* |
| 1 | 2 | 4 | 0.156654 | 0.081497 | 0.162688 | 0.099161 | -0.0091 | 0.159813 |
| 1 | 2 | 5 | 0.197112 | 0.082379 | 0.109589 | 0.11092 | 0.051343 | 5.13317* |
| 1 | 2 | 6 | 0.215739 | 0.072776 | 0.123004 | 0.088481 | 0.040548 | 4.01347* |
| 1 | 3 | 4 | 0.124449 | 0.091552 | 0.176875 | 0.107124 | 0.011447 | 0.251911 |
| 1 | 3 | 5 | 0.236392 | 0.049819 | 0.118733 | 0.095055 | 0.066221 | 11.7868* |
| 1 | 3 | 6 | 0.176321 | 0.05693 | 0.167597 | 0.099152 | 0.031765 | 2.017614 |
| 1 | 4 | 5 | 0.186545 | 0.095081 | 0.071434 | 0.14694 | 0.082476 | 10.5217* |
| 1 | 4 | 6 | 0.180297 | 0.096093 | 0.109022 | 0.114587 | 0.040734 | 2.888049 |
| 1 | 5 | 6 | 0.172502 | 0.079552 | 0.149065 | 0.098881 | 0.020795 | 0.808814 |
| 2 | 3 | 4 | 0.143189 | 0.087697 | 0.173774 | 0.09534 | 0.006351 | 0.079209 |
| 2 | 3 | 5 | 0.239605 | 0.069337 | 0.103666 | 0.087392 | 0.055007 | 8.41204* |
| 2 | 3 | 6 | 0.141078 | 0.084521 | 0.169385 | 0.105016 | -0.002 | 0.007681 |
| 2 | 4 | 5 | 0.201763 | 0.110214 | 0.069613 | 0.11841 | 0.064874 | 8.07710* |
| 2 | 4 | 6 | 0.074161 | 0.142371 | 0.142371 | 0.141097 | 0 | 0 |
| 2 | 5 | 6 | 0.172724 | 0.088155 | 0.142527 | 0.096595 | 0.016479 | 0.511758 |
| 3 | 4 | 5 | 0.157883 | 0.115863 | 0.089558 | 0.136697 | 0.044822 | 2.926938 |
| 3 | 4 | 6 | 0.098566 | 0.141188 | 0.131539 | 0.128707 | 0.023542 | 0.76588 |
| 3 | 5 | 6 | 0.15671 | 0.10181 | 0.153616 | 0.087864 | -0.00748 | 0.101766 |
| 4 | 5 | 6 | 0.137416 | 0.101387 | 0.156667 | 0.10453 | 0.00608 | 0.062021 |

\*P-value <0.05. a: The data came from MAPMAKER/EXP(3.0b)[27]

**Table 2.** The ELS estimated frequencies of nonsister gametes in triplets of 6 dominant loci in 333 F2 mice^a^ and recombination disequilibrium tests

| Locus | | | $p_1 =$ | $p_2 =$ | $p_3 =$ | $p_4 =$ | RD | |
| --- | --- | --- | --- | --- | --- | --- | --- | --- |
| a | b | c | $p(abc)$ | $p(Abc)$ | $p(abC)$ | $p(aBc)$ | $D_r$ | $\chi^2$ |
| 1 | 2 | 3 | 0.208668 | 0.086162 | 0.094698 | 0.110472 | 0.059571 | 4.19998* |
| 1 | 2 | 4 | 0.14751 | 0.087958 | 0.160866 | 0.103665 | 0.004568 | 0.038389 |
| 1 | 2 | 5 | 0.200976 | 0.080676 | 0.108494 | 0.109854 | 0.053301 | 3.454827 |
| 1 | 2 | 6 | 0.192229 | 0.084408 | 0.126033 | 0.09733 | 0.032286 | 1.529476 |
| 1 | 3 | 4 | 0.140566 | 0.079252 | 0.181795 | 0.098387 | -0.00231 | 0.011286 |
| 1 | 3 | 5 | 0.209093 | 0.065783 | 0.12237 | 0.102753 | 0.05374 | 3.709605 |
| 1 | 3 | 6 | 0.16895 | 0.063323 | 0.165648 | 0.102079 | 0.027028 | 1.052238 |
| 1 | 4 | 5 | 0.173539 | 0.100396 | 0.079665 | 0.1464 | 0.069631 | 4.37455* |
| 1 | 4 | 6 | 0.16482 | 0.102771 | 0.113819 | 0.118591 | 0.031395 | 1.203758 |
| 1 | 5 | 6 | 0.173447 | 0.07932 | 0.148645 | 0.098588 | 0.021237 | 0.683502 |
| 2 | 3 | 4 | 0.141958 | 0.088919 | 0.172784 | 0.096339 | -0.00675 | 0.088422 |
| 2 | 3 | 5 | 0.202566 | 0.085775 | 0.112098 | 0.099561 | 0.04221 | 2.458404 |
| 2 | 3 | 6 | 0.139672 | 0.086032 | 0.16843 | 0.105866 | 0.001185 | 0.002673 |
| 2 | 4 | 5 | 0.173634 | 0.117152 | 0.085607 | 0.123607 | 0.045733 | 2.356028 |
| 2 | 4 | 6 | 0.10113 | 0.133668 | 0.133668 | 0.131535 | 0 | 0 |
| 2 | 5 | 6 | 0.156859 | 0.096648 | 0.142758 | 0.103735 | 0.009898 | 0.150093 |
| 3 | 4 | 5 | 0.143582 | 0.120278 | 0.098137 | 0.138002 | 0.032044 | 1.109958 |
| 3 | 4 | 6 | 0.105025 | 0.139707 | 0.129181 | 0.126086 | 0.019221 | 0.440547 |
| 3 | 5 | 6 | 0.140841 | 0.109125 | 0.152351 | 0.097682 | -0.01147 | 0.211908 |
| 4 | 5 | 6 | 0.146847 | 0.096392 | 0.156905 | 0.099856 | -0.00184 | 0.005869 |

\*P-value <0.05. a: The data came from MAPMAKER/EXP(3.0b)[27]

**Table 3.**
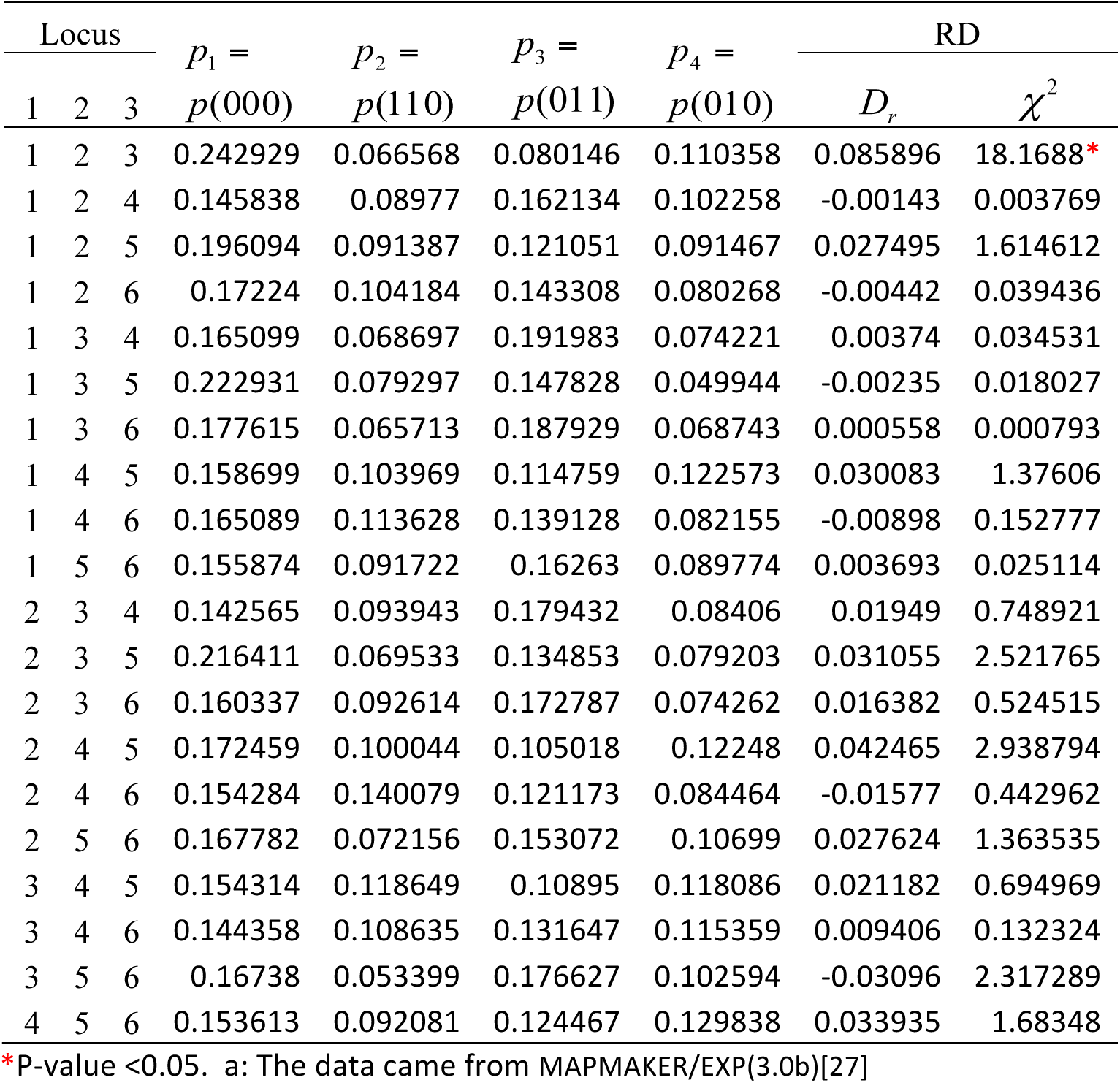
The moment estimated frequencies of nonsister gametes in triplets of 6 codominant loci in 333 F2 mice^a^ and recombination disequilibrium tests

### Test for recombination disequilibrium among three genes in testcross phenotype data

A well-known example for testing for crossover interference among three genes in testcross/backcross phenotype data is the female testcross 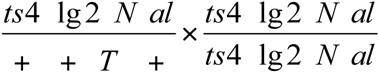 for the 3L.34 breakpoint in maize where *T* and *N* denote translocation breakpoint and normal point, respectively. The data can be found in Auger and Sheridan (Auger and Sheridan 2001) and Tan and Fornage (Tan and Fornage 2008). In this testcross, the single crossover within interval II (*lg*2-*T*) did not occur but double crossovers between adjacent intervals II (*lg*2-*T*) and III (*T-al*) did occur. The recombination fractions within intervals II (*lg2-T*) and III (*T-al*) are *r_lg_*_2_*_T_* = 0+0.0047=0.0047 and *r_Tal_* = 0.2025+ 0.0047= 0.2072 and frequency of the observed double crossover types is *r_DC_* = 0.0047. RD among the three loci (*lg2-T-al*) is *D_r_* = *r_DC_* − *r_lg_*_2_*_T_r_Tal_* = 0.00372, which has 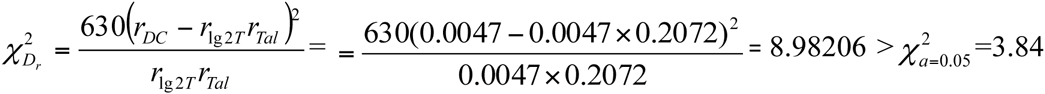 at p-value <0.05 level. These two examples have a common point that at least one of two adjacent intervals is very short (Tan and Fornage 2008; Tan and Fu 2007).

Another famous example also comes from *Drosophila*. There we have *v_cv*^+^*_ct*^+^ (parental type): 580; *v*^+^*_cv*_ct (parental type): 592; *v*^+^*_cv_ct*^+^ (single crossover between genes *ct* and *cv*): 45; *v_cv^+^_ct* (single-crossover between genes *ct* and *cv*): 40; *v_cv_ct* (single crossover between genes *v* and *ct*): 89; *v*^+^*-cv*^+^-*ct*^+^ (single-crossover between genes *v* and *ct*): 94; *v*^+^*_cv*^+^*_ct* (double-crossover): 3; and *v_cv_ct*^+^ (double crossover): 5. Total number of gametes is 1448 (Anthony *et al*. 2008). From this data, the frequencies of the four types of nonsister gametes are estimated as

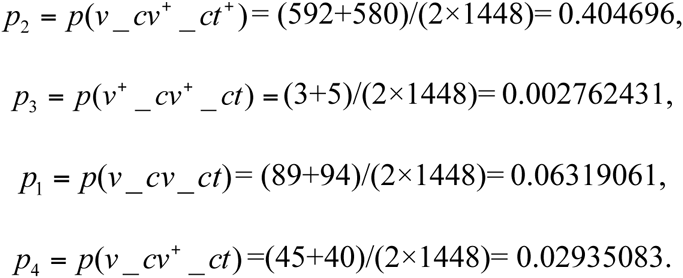

Thus, RD among the three genes *v*, *cv*, and *ct* is *D_r_* = *4*(*p*_2_*p*_3_ − *p*_1_*p*_4_) = ™0.0029470 and 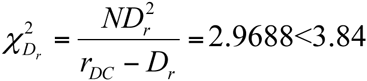. Therefore, this positive interference among the three genes is due to experimental error.

### Evaluation of genetic map

An important application of RD is to evaluate genetic map of a chromosome or a chromosome fragment. If a chromosome map is correct and the three-locus RD is significant, then the three loci must be located within an adjacent region. Otherwise, some markers of this map would be falsely linked because a significant RD implicates a strong recombination interference that occurs between two adjacent shorter intervals. To elucidate this point, we here chose a genetic map of rice chromosome 12 made by Harushima et al (Harushima *et al*. 1998). This chromosome map (here is called Harushima rice map 12 for convenience) consisting of 93 loci was built with126 markers in an F_2_ population, among which 90 are codominant and 36 are dominant. For convenience, we here just considered the 90 codominant markers and used the raw data (downloaded from the website: http://rgp.dna.affrc.go.ip/public) to construct 117481 triplets. 4 non-sister gamete frequencies *p*_1_, *p*_2_, *p*_3_, and *p*_4_ in each triplet were estimated by applying our method(Supplementary Note S2). The results are summarized in Supplementary Table1 (excel sheet1). By testing for RDs using Equations (6) (since there are strong linkage relationships among loci and each marker locus repeatedly occurs in many triplets, Bonferroni or Benjamini-Hochberg multiple tests are not available to test the 117481 RDs), we found 137 triplets with significant RDs at significant level of 0.05. Supplementary Table 2S (excel sheet2) shows that these 137 triplets form at least 12 larger linkage triplet groups: G24B, V57B, R3266, F8, S790B, C362A, C1116A, R769B, S2572, G193, R2253B, and C732B. Linkage triplet group G24B consists of 8 triplets (G24B, R2253B, C930), (G24B, S10637A, S10363), (G24B, V110, S1830), (G24B, R1709, R2672B), (G24B, C87, R496), (G24B, S861, R2672B), (G24B, L714, C930), (G24B, R1759, S13561). Among them, markers R2253B, V110, L714, and S10637A have their own linkage triplet groups, markers S10363, S1830, R1709, R2672B, C87, R496,S861, L714, C930, R1759, and S13561 are separately scattered in triplet groups V57B, R3266, V9A, R2253B, R642B, S894, R769B, S2572, V9A, S790B, R1957, F8, V110, C1116A. However, these markers were mapped into an end (72.9cM ~ 109.3cM) of Harushima rice map 12, while G24B was mapped into another end (5.5 cM) (see Fig. 1A). On the other hand, the markers V57B, R3266, F8, G193, R2253B, C732B, S790B, R3025S, C362A, C104A, G1112, R328A, W120B, R1957, and C1116A that are tightly linked to G24B on the rice map12 are not members of G24B triplet group. Again, markers R496, R1759, S1830, G1106, R2292, C930, S13561, and L405B were located at the same position (108.8 cM) on the Harushima rice map 12, but except for C930, S1830, R496, and S13561 that are members of G24B and V57B linkage triplet groups, all the others are scattered in different linkage triplet groups, indicating that it is impossible that these 8 markers were located at the same linkage position. Markers V124, S11679, M10C, and T5 were also located at the same linkage position of 70.7cM, but they are not found in these 137 linkage triplets with significant RD. To globally display RD chi-square tests of 137 triplets, we constructed a 4D plot (Fig. 1B). To easily see incorrect of the Harushima rice map 12 (Fig 1.A), we gave the RDs expected to occur between two adjacent intervals among three adjacent loci mapped in the Harushima rice map 12. The expected RDs follow on red diagonal dots. Theoretically, if the Harushima rice map 12 is correct, then RDs should distribute along the red diagonal dots between original point (0,0,0) and end point (90,90,90) and in its neighboring regions. However, Fig. 1B shows that almost all of these 137 RD points are far away from this diagonal line. These indicate that Harushima et al’s map 12 is not correct.

**Figure 1.**
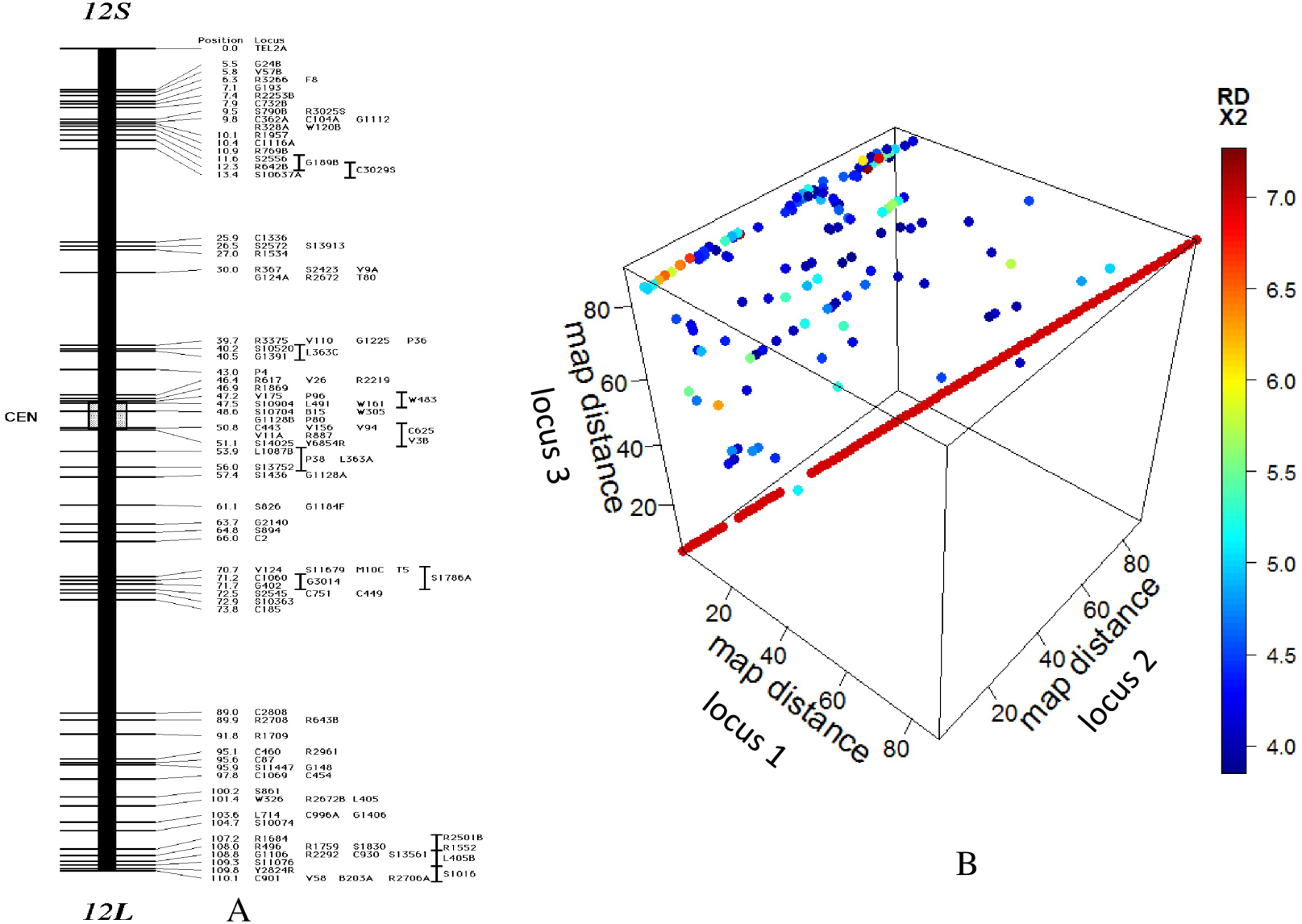
Genetic map of rice chromosome 12 and 4D RD plot. A: Genetic map of rice chromosome 12 was made with 126 markers among which 90 are codominant markers and 36 are dominant makers. B: RDs of 137 triplets that were detected to be significant from 117481 triplets constructed with 90 codominant markers from rice chromosome 12 were plotted in 4Dplot(3D-space for 90 markers and 1D color for RD values). Diagonal red dots from (0,0,0) to (90,90,90) are theoretical RD among three adjacent loci.

### Haplotypes

Most of SNP haplotype data were derived from human populations that are not ideal population. To demonstrate that the RD test can be also applied to natural populations, we here used a haplotype dataset published by Fallin et al (Fallin *et al*. 2001) as an example of RD analysis in natural populations. The haplotypes consist of 4 SNPs scattered in a 205kbp region on human chromosome 19. The map of the four SNP loci is M1M3M4M6. In the results of Fallin et al, no LD was found between M1 and M4 and between M1 and M6 but LD existed between M4 and M6, between M3 and M4, and between M3 and M6 with p-value<0.001, suggesting that M3, M4 and M6 are in short region. To perform triplet RD tests, we constructed four combinations of three-SNP haplotypes: M1M3M4, M1M4M6, M3M4M6 and M1M3M6 each having 8 haplotypes. The results of our RD analysis of four combinations of three-haplotypes were summarized in Table 4. Table 4 shows that there are no RDs among M1, M4 and M6, among M1, M3 and M6, while M3, M4 and M6 had very significant RD, which is very consistent with significant LDs between M3 and M4, between M4 and M6 and between M3 and M6.

**Table 4.** RD and chi-square testing RD among three SNPs in four haplotypes (M1M3M4*M6) where M4* is C19M4 that is part of ApoE-ε4

|  | M1M3M4* | M1M4*M6 | M3M4*M6 | M1M3M6 |
| --- | --- | --- | --- | --- |
| **P1** | 0.475 | 0.305 | 0.589 | 0.32 |
| **P2** | 0.032 | 0.199 | 0.358 | 0.175 |
| **P3** | 0.472 | 0.292 | 0.008 | 0.316 |
| **P4** | 0.023 | 0.206 | 0.047 | 0.191 |
| **RD** | -0.0042 | 0.0047 | 0.0248 | 0.0058 |
| **X<sup>2</sup>** | 0.237 | 0.041 | 10.247 | 0.067 |
| **p-value** | 0.626 | 0.839 | 0.0014 | 0.795 |

## Discussion

Estimation of gamete frequencies is the first step to test for RD among three or multiple loci. Therefore a good method for accurately estimating gamete frequencies in a population is required. In F_2_ population, for dominant loci, the current existing EM methods have low power (Liu 1998) because these EM methods still inefficiently distinguish dominant homozygous genotypes from dominant heterozygous genotypes and have to use a lot of trivial and complex algebraic equations to overcome this difficulty, while the TF method efficiently utilizes sister gamete genotypes with equal frequencies to reduce complexity of estimation of gametic frequencies. The modified TF method tries to find a desirable value of *q*_1_ such that difference between the observed and expected frequencies of gametes is relatively small. Therefore, it can give more accurate estimation of gamete frequencies than the TF method. This point was supported by the above results of testing for RD among loci obtained from the real genotype data (Tables 1–3). For codominant loci, the EM algorithms have really high power to estimate frequencies of three-locus gametes (Li et al 2005 and Esch 2005), however, the computational burden would be extremely huge when number (n) of loci is very large because there would be n^3^ triplet gametes (Esch 2005). For example, in general GWAS data, number of SNPs is more than 2000000, so there would be 2000000^3^ = 8 × 10^18^ three-locus gametes. To significantly reduce computational burden, estimation of gamete frequencies of genome-wide loci requires both fast and powerful method. This is why here we want to propose this new method. Since all genotypes can be recognized and are informative for estimation of gamete frequencies, there is not phase problem to be solved. In addition, likewise, the assumption that sister gametes also have the same frequencies reduces complexity of estimation equations. For example, we just use 10 simple binomial equations to estimate frequencies of four non-sister gametes, while Li et al (2005)’s EM method requires 18 complex algebraic equations and Hospital et al (Hospital *et al*. 1996)’ method requires 20 equations. In natural or outbreeding populations, frequencies of gametes or haplotypes can be estimated by using the existing powerful methods such as PHASE(Stephens *et al*. 2001), fastPHASE(Scheet and Stephens 2006), BEAGLE (Browning and Browning 2007), IMPUTE2 (Howie *et al*. 2009) and MaCH (Li *et al*. 2010).

One application of RD is the construction of genetic maps for fine mapping. As seen in Tables 1–3, some triplets have significant RD, but most of triplets have no significant RD. For the triplets without significant RD, their coincident coefficients are set to be 1; for those with significant negative or positive RD, the coincident coefficients are given by Tan and Fornage definition (Tan and Fornage 2008). Thus, one can use Tan and Fornage mapping functions (Tan and Fornage 2008) to accurately calculate the genetic distances between these markers.

Since RD provides information of recombination interference among three loci, it is useful for the evaluation of the linkage map. As seen above, Harushima rice map12 is very inconsistent with results of our triplet analysis. One possible reason is that this map was based on two-point analysis and the two-point approach does not utilize information of recombination interference occurring between two adjacent intervals, while recombination interference disturbs linkage information between loci. As a result, two-point analysis would generate incorrect linkage between loci. Using three-locus EM methods or our methods (Supplementary Note S2), one can obtain recombination fractions among three adjacent loci, and then use the method provided by Tan and Fu (Tan and Fu 2007) to convert three-locus recombination fractions into two-locus recombination fractions. A correct genetic map can be made by using UG mapping method (Tan and Fu 2006) or other mapping methods, the chi-square test for the three-locus RD and Tan-Fornage map functions (Tan and Fornage 2008).

In ideal population such as F_2_ or backcross population, RD is purely due to recombination interference between short intervals but in natural populations, in addition to recombination interference, RD may also result from selection, mutation, drift, and gene conversion because these evolutionary factors also change frequencies of gametes. So as with LD, evolutionary history of multiple genes can be revealed by testing for RD among genes. In addition, RD can also be applied to genome-wide association analysis of haplotypes with diseases. For this application, we have furthermore developed a new method for studying association of haplotypes with disease of study and in somewhere we will publish it.

